# uORFlight: a vehicle towards uORF-mediated translational regulation mechanisms in eukaryotes

**DOI:** 10.1101/713321

**Authors:** Ruixia Niu, Yulu Zhou, Rui Mou, Zhijuan Tang, Zhao Wang, Guilong Zhou, Sibin Guo, Meng Yuan, Guoyong Xu

**Affiliations:** State Key Laboratory of Hybrid Rice, Institute for Advanced Studies (IAS), Wuhan University, Wuhan, Hubei 430072, China; Guangxi Key Laboratory of Rice Genetics and Breeding, Rice Research Institute, Guangxi Academy of Agricultural Science, Nanning, Guangxi 530007, China; National Key Laboratory of Crop Genetic Improvement, National Centre of Plant Gene Research (Wuhan), Huazhong Agricultural University, Wuhan, Hubei 430070, China

## Abstract

Upstream open reading frames (uORFs) are prevalent in eukaryotic mRNAs. They act as a translational control element for precisely tuning the expression of the downstream major open reading frame (mORF) with essential cellular functionalities. uORF variation has been clearly associated with several human diseases. In contrast, natural uORF variants in plants have not ever been identified or linked with any phenotypic changes. The paucity of such evidence encouraged us to generate this database-uORFlight (http://uorflight.whu.edu.cn). It facilitates the exploration of uORF variation among different splicing models of Arabidopsis and rice genes. Most importantly, users can evaluate uORF frequency among different accessions at the population scale and find out the causal single nucleotide polymorphism (SNP) or insertion/deletion (INDEL) which can be associated with phenotypic variation through database mining or simple experiments. Such information will help to make hypotheses of uORF function in plant development or adaption to changing environments on the basis of the cognate mORF function. This database also curates plant uORF relevant literature into distinct groups. To be broadly interesting, our database expands uORF annotation into more species of fungi (*Botrytis cinerea*), plant (*Brassica napus, Glycine max, Gossypium raimondii, Medicago truncatula, Solanum lycopersicum, Solanum tuberosum, Triticum aestivum* and *Zea mays*), metazoan (*Caenorhabditis elegans* and *Drosophila melanogaster*) and vertebrate (*Homo sapiens, Mus musculus* and *Danio rerio*). Therefore, uORFlight will light up the runway toward how uORF genetic variation determines phenotypic diversity and advance our understanding of translational control mechanisms.

## Introduction

Gene expression must be tightly regulated at transcription, translation and post-translation levels. The imperfect correlation between protein abundance and mRNA levels suggests translational efficiency regulated by translational control as one of the determinants of protein output from variable mRNA input. This layer of regulation is mediated by the cooperative action between different mRNA elements and trans-acting factors (1). Upstream open reading frames (uORFs) are among the mRNA elements that can confer precise control of protein translation.

A uORF initiation codon resides upstream of the coherent mORF, and will be first encountered by 43S scanning ribosome (including 40S ribosomal subunit and eIF2 ternary complex). Sequentially, 60S subunit joins in and reconstitutes 80S ribosome for uORF translation elongation, after which the 40S and 60S are disjointed and 40S may remain associated with mRNA. Therefore, usually uORF translation is prioritized over mORF, leading to hindered translation of the mORF. Only in situations where the remaining 40S ribosomes regain fresh eIF2 ternary complex and other unknown reinitiation factors, or when uORF initiation codon is bypassed by the scanning ribosome, the downstream mORF has the chance to be translated (Figure 1a). The former situation is termed as reinitiation and the latter as leaky scanning, two mechanisms that have been accepted as an explanation of limited mORF expression under normal growth and developmental conditions (2-4). Most importantly, a uORF can confer selective mORF translation in response to a wide-range of cellular stimuli, such as metabolite and ion homeostasis, hormone changes, environmental signals, and immune induction (See uORF references on this website). This tight and temporal regulation pattern fine-tunes the translational efficiency of mORFs and thus guarantees appropriate protein quantity and quality for adaption to different physiological conditions. Because of those unique features, we have successfully utilized uORF-mediated translational control in engineering disease resistant plants without fitness costs by restricting toxic resistance protein translation under normal conditions but allowing transient induction under pathogen infection conditions (5).

**Figure 1.**
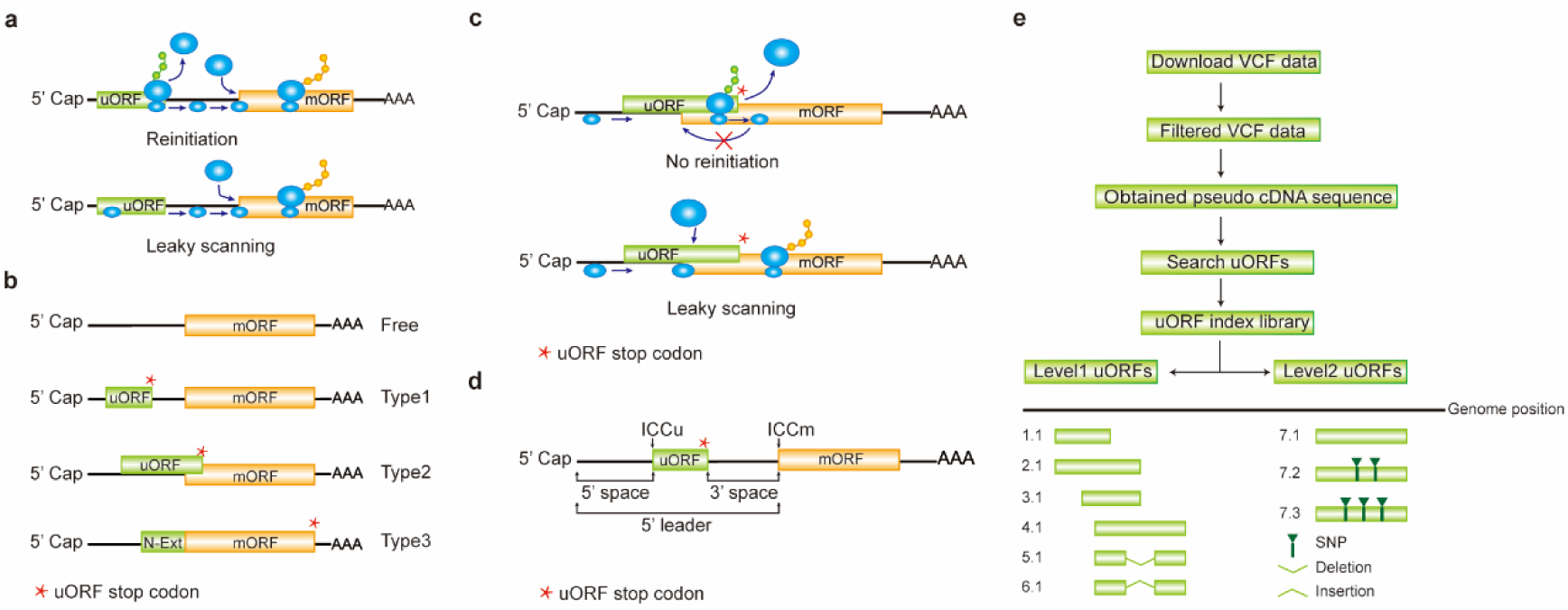
Computation of uORF variation. **a**, Reinitiation and leaky scanning models. In the reinitiation model, the 80S ribosomal subunit will separate after translating the uORF and the 40S subunit remains associated with mRNA, regaining fresh eIF2 ternary complex and other unknown reinitiation factors to translate the mORF. In the leaky scanning model, the uORF initiation codon is bypassed by the scanning complex, which will ignore the uORF and translate the mORF. **b**, uORF types. Genes without uORFs are grouped as ‘Free’. uORF are further divided into Types1-3 with respect to the position of uORF stop codon relative to the mORF. N-Ext, N extension. **c**, Type2 uORF-controlled mORF translation is only favored by leaky scanning. Overlap between the Type2 uORF and mORF makes reinitiation of the mORF after translation of the uORF impossible. **d**, uORF positional information on cDNA. The mORF is flanked by the 5’ leader and 3’ UTR (3’ untranslated region). 5’ and 3’ space are used to describe the distance from Cap to uORF AUG and from uORF stop codon to mORF AUG, respectively. The sequence from −3 to +4 relative to the AUG initiation codon (A as +1) corresponding to the Kozak consensus (A/GCCAUGG) position is termed as initiation codon context (ICC) with ICCu and ICCm for uORF and mORF, respectively. **e**, Workflow to identify uORF variants.

However, once uORF-mediated precision control has been challenged by genetic variation or mis-regulation, it causes human diseases (6, 7). By 2009, 509 human genes had been identified with polymorphic uORFs and some of them have been experimentally associated with different human diseases, including malignancies, metabolic or neurologic disorders, and inherited syndromes. This trend became more striking recently with more genomic variation data released and analyzed (3, 8). In contrast, natural variation of plant uORFs has not yet been investigated, even though there are abundant publicly accessible genetic and phenotypic variation data, especially for Arabidopsis (*Arabidopsis thaliana*) and rice (*Oryza sativa* L.). Since the release of the Arabidopsis reference genome of accession Col-0 in 2000, the rapid development of sequencing technology has bolstered genome-wide association studies (GWAS) by linkage disequilibrium of interesting phenotypic traits with the most probable genetic variation, particularly using the genome sequences of 1135 accessions from the 1001 Genomes Project (9-15). Genetic variation of rice has long been used for molecular breeding and recent re-sequencing of a large set of rice accessions, especially from the 3000 Rice Genomes Project, generated a wealth of genetic variation for the discovery of useful alleles for agronomic trait improvement (16-24).

It is noteworthy that previous genotype-phenotype association studies mainly focused on protein coding regions, while it is becoming more evident that the cis-element variation, such as in the promoter regions, weighs a lot in determining phenotypic variation (25). However, there is still less attention that has been paid on the variation of mRNA regulatory elements, such as uORFs. In this study, we used public resources to identify uORF variation for further experimental verification of phenotypic diversity mediated by translational control.

## Methods

A uORF is defined as the presence of an initiation codon in an annotated mRNA 5’ leader region and can be categorized into ‘Types 1-3’ based on the positions of uORF stop codon, with Type1 in 5’ leader, Type2 in mORF coding region and Type3 shared with mORF (also known as an mORF N-extension). We term those mRNAs without a uORF as ‘Free’. It is obvious that reinitiation is impossible for translation of Type2 uORF-controlled mORFs (Figure 1b and c). Hereafter, ORF of both uORF and mORF means that AUG is used as the initiation codon, unless specifically stated. We use the term 5’ leader sequence instead of 5’ untranslated region (5’ UTR), considering the peptide-coding potential of uORFs (26, 27). The sequence from −3 to +4 relative to AUG initiation codon (A as +1) corresponding to Kozak consensus (A/GCCAUGG) position is termed as initiation codon context (ICC) with ICCu and ICCm for uORF and mORF, respectively (Figure 1d).

We chose the Arabidopsis Col-0 accession (Ensemble V39; TAIR11) and rice Nipponbare cultivar (MSU V7) as reference genomes for dicot and monocot uORF analysis, respectively. Arabidopsis representative gene models (27445 in total; Supplementary Table 1 in **Download** menu) of the nuclear coding proteins contain 26713 genes from TAIR10 representative gene annotation file and 732 genes using their ‘.1’ splicing models. Nipponbare representative gene models (38860 in total; Non-TE Loci) are defined using their smallest numbered models (38618, 221, 16 and 5 genes using the ‘.1’, ‘.2’, ‘.3’ and ‘.4’ splicing models, respectively; Supplementary Table 2 in **Download** menu). To calculate uORF variation, we downloaded VCF (variant call format) files of Arabidopsis 1135 accessions from the 1001 Genomes Project and rice 3k varieties from the 3000 Rice Genomes Project (9, 20, 24, 28). We filtered SNPs and INDELs with low quality indicated in the VCF files and used the alleles with frequency over 90% as suggested (28). We further removed genes with variants affecting the annotated initiation codon of their mORF.

We replaced all the splicing models with filtered variants and searched uORFs of all the accessions. We then transferred the relative positions of uORFs on the different transcripts into absolute positions in the referenced genome and assigned continuous index numbers starting with number 1.1 in the order of uORF ATG occurrence in the genome. Those uORFs with the same genomic positions and sequences are assigned the same index number to indicate no uORF changes. Otherwise, two leveled uORF identifiers (Level1.Level2) were used to describe the variation. Level1 indicates major differences that cause uORF creation, loss or length changes due to SNPs or INDELs. Level2 indicates minor differences that lead to nucleotide and/or amino acid substitution due to SNPs (Figure 1e). We grouped Arabidopsis accessions on the basis of the latitude at which they were collected (15-degree interval; Supplementary Table 1 in **Download** menu). The frequency of an individual uORF in Arabidopsis was calculated based on its occurrence in the total population and in different latitude ranges. The frequency in rice was calculated as its occurrence in total population and nine subgroups (20). The associated SNP or INDEL identifiers are also recorded along with uORF variants.

MySQL database schema was used for uORF information storage and a user-friendly PHP web interface was designed to query and download. GO analysis was done using Omicshare online tools (http://www.omicshare.com/) with the default setting. The uORF annotation information of the other species can be found in the website.

## Results and Discussion

With the recent recognition of the significance of uORFs within distinct physiological contexts, the following functionalities will help the community quickly overview the progress in this area and find out uORF variation to link with phenotypic diversity at the population level.

### uORF in the reference genomes

uORFs are becoming increasingly attractive because of their capacity to fine-tune translational control and respond accurately to distinct extracellular and intracellular stimuli. However, the current understanding of uORFs is based on a small number of case studies. In an attempt to provide guidelines, we investigated natural patterns in uORF types, length distribution and ICC of Arabidopsis and rice representative gene models. uORFs are more prevalent in Arabidopsis (48.45%), and the lower frequency (20.65%) of uORFs in rice transcripts may arise from incomplete 5’ leader annotation. Their prevalence is mostly due to overrepresentation of Type1 uORFs which account for 90.79% and 87.63%, in contrast to only 9.16% and 12.17% for Type2 uORFs of Arabidopsis and rice, respectively. The Type3 uORFs are the least common (19 uORFs in 17 Arabidopsis genes, and 74 uORFs in 41 rice genes), perhaps because they may give rise to N-extension and are likely to alter protein activities or molecular localizations as reported (29). Type2-containing genes tend to occur along with Type1 uORFs, and the significance of their co-existence needs further investigation (Figure 2a). GO analysis shows clearly different enriched terms among those uORF-Free, Type1-only and Type2 genes in both Arabidopsis and rice, suggesting that uORF type attributes could have an impact on different functional groups of genes (Figure 2b; Supplementary Table1 and 2 in **Download** menu). This assumption appears to be reasonable because both leaky scanning and reinitiation mechanisms could overcome Type1 uORF inhibition, while only leaky scanning can overcome Type2 uORF inhibition (Figure 1a and b), and leaky scanning seems to be of low efficiency in both animals and plants (30, 31). Importantly, uORF-containing genes are enriched in more specific groups than uORF-Free genes. Moreover, even though rice has fewer enriched GO-terms, most of them (89.47%) are shared with GO-terms enriched for Arabidopsis uORF-containing genes. The 11 shared GO-terms for Type1 only and Type2 include ‘signaling’ and ‘cellular response to stimulus’ (Figure 2c). All these results suggest that uORFs are more frequently used by specific functional groups and these groups are shared between Arabidopsis and rice. Next, we examined uORF length distribution and found that Type2 uORFs are on average longer than Type1, and that rice uORFs are generally longer than those of Arabidopsis (Figure 2d). A minimal Type1 uORF consisting of just an AUG and a stop codon has been shown to be sufficient for translational inhibition of three boron (B)-related genes and also sufficient for translational responsiveness to low B stimulation (32). Such short uORFs (6.79%) are more commonly found among Arabidopsis Type1 uORFs and must require other synergistic cis-elements to allow translational responsiveness to specific stimuli. Therefore, the attribute of uORF length appears not to be a definitive parameter for prediction of uORF function. We then asked whether uORF surrounding sequences display some informative patterns. We found that only 0.45% ICCu and 2.99% ICCm in Arabidopsis and 1.93% ICCu and 11.88% ICCm in rice contain Kozak consensus sequences, suggesting that more variable contexts are used to flexibly tune ORF translation in nature. Enrichment analysis of ICCu and ICCm did not uncover any obvious enriched sequence, suggesting the feasibility of engineering tailored protein expression using ICC variants in plants.

**Figure 2.**
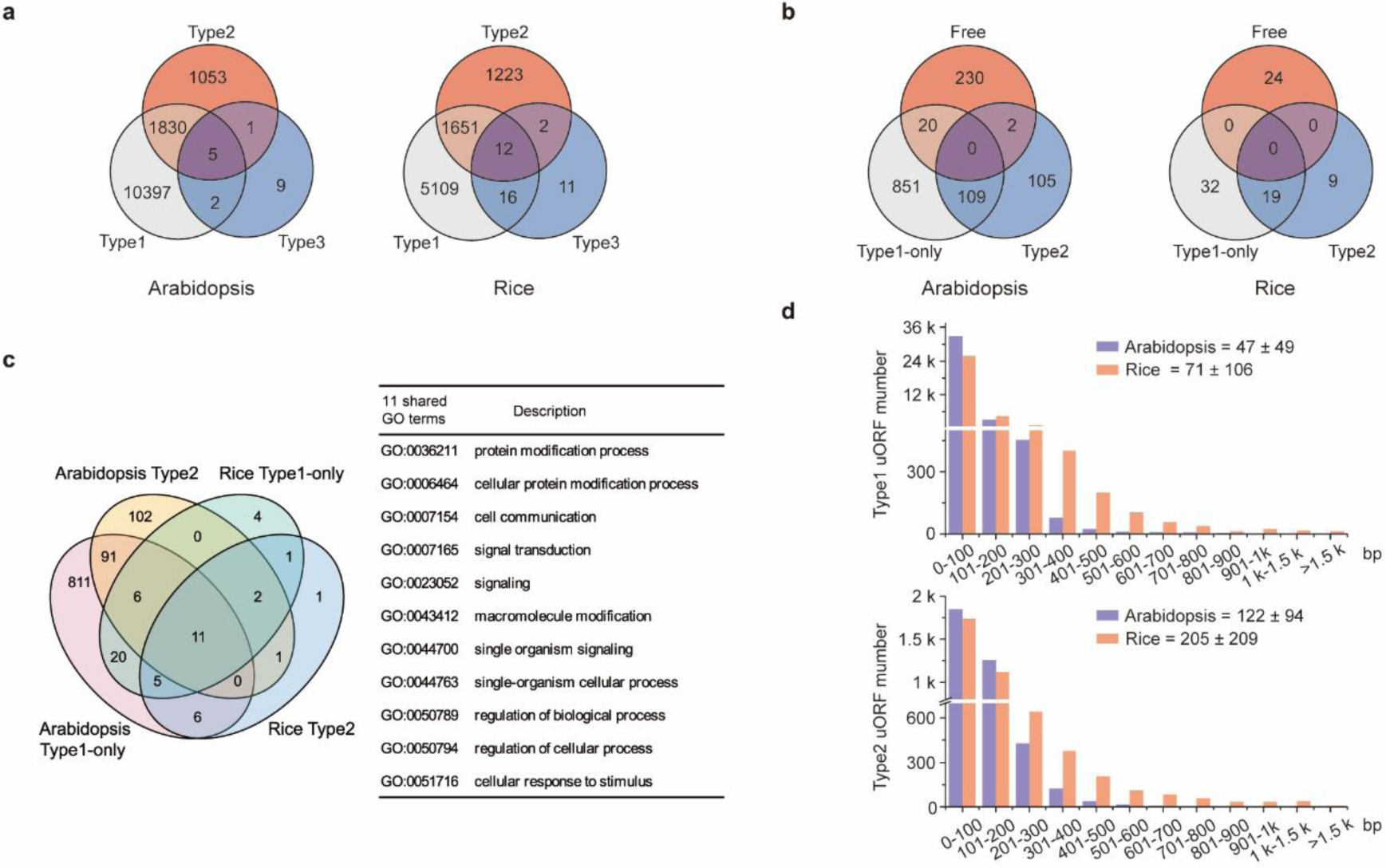
Characterization of uORFs in Arabidopsis and rice. **a**, Venn diagram of uORF type distribution among uORF-containing genes. **b**, Venn diagram of enriched Gene Ontology (GO) terms of uORF-Free-, Type1-only- and Type2-uORF-containing genes in Arabidopsis and rice. GO terms of FDR (false discovery rate) < 0.05 are used. **c**, Venn diagram to show GO-terms shared in Arabidopsis and rice uORF-containing genes. 51 out of 57 enriched GO-terms of rice uORF-containing genes are also found in those of Arabidopsis. 11 GO-terms enriched in both Type1-only and Type2 uORF-containing genes of Arabidopsis and rice are detailed in the table. **d**, Length distribution of Type1 and Type2 uORFs. Insert number shows Mean±SD of uORF length (bp). The representative gene models of Arabidopsis reference accession Col-0 and rice reference cultivar Nipponbare are used for analysis.

From **uORF view** menu, users can browse and search uORF information for all splicing models in the reference genomes, including eudicot Arabidopsis Col-0 and monocot rice Nipponbare. The entirety of the uORF information is also saved in a table format (Supplementary Table 1 and 2 in **Download** menu) which can be accessed in the ‘Download’ Menu. Choosing the reference genome will pop out a search page, this page requires users to input the splicing model identifier (ID; e.g. AT4G36990.1) and the search results will return the summary information of the uORF, details of which can be accessed by clicking the ID or downloaded in the top-right corner in the popped out page. uORF annotation of other species can be obtained in a similar way under this menu.

### uORF variation at the population level

uORF variation has been studied at the genome-wide scale using human SNPs and the results provide clear evidence of its association with genetic diseases. However, these studies only assessed single SNP effects that result in uORF initiation codon creation or stop codon loss (3, 8). We analyzed uORF variation by considering the integral effect of all homozygous SNPs and INDELs in each accession. Significantly, we found that 54.17% of Arabidopsis uORF variants (64.52% in rice) of the representative gene models are associated with at least two genetic changes. As emphasized above, uORF creation, loss, length changes, and type switches are considered as major variation and are defined as Level1 (Figure 1e). Level2 variation is likely to affect uORF function, provided that its nucleotide sequence or encoded peptide is able to cause ribosome stalling or mRNA decay (4, 6, 7). To simply show the significance of uORF variation, we focused on three Level1 variants which alter uORF type attributes between Type1-only and Type2, between uORF-Free and Type1-only, between uORF-Free and Type2, which excludes Level1 changes (Figure 3a-c). We first found that 31.03% and 20.40% of uORF-containing genes have altered uORF type attributes among different splicing models in Col-0 and Nipponbare, respectively, suggesting that uORF variation produced by alternative splicing may add translational control regulation to specific processes (Figure 3c and d; Supplementary Table 3 and 4 in **Download** menu). We also identified the three Level1 variants of different accessions at the population level by firstly analyzing the representative gene models (Figure 3e; Supplementary Table 3 and 4 in **Download** menu). Alternatively, we identified the three Level1 variants of different accessions at the population level by considering all splicing models of a defined gene, and any splicing model of one accession different from the other accession are counted (Figure 3f; Supplementary Table 3 and 4 in **Download** menu). Significantly, these two different ways exemplify the importance of uORF variation on key genes of different cellular functions.

**Figure 3.**
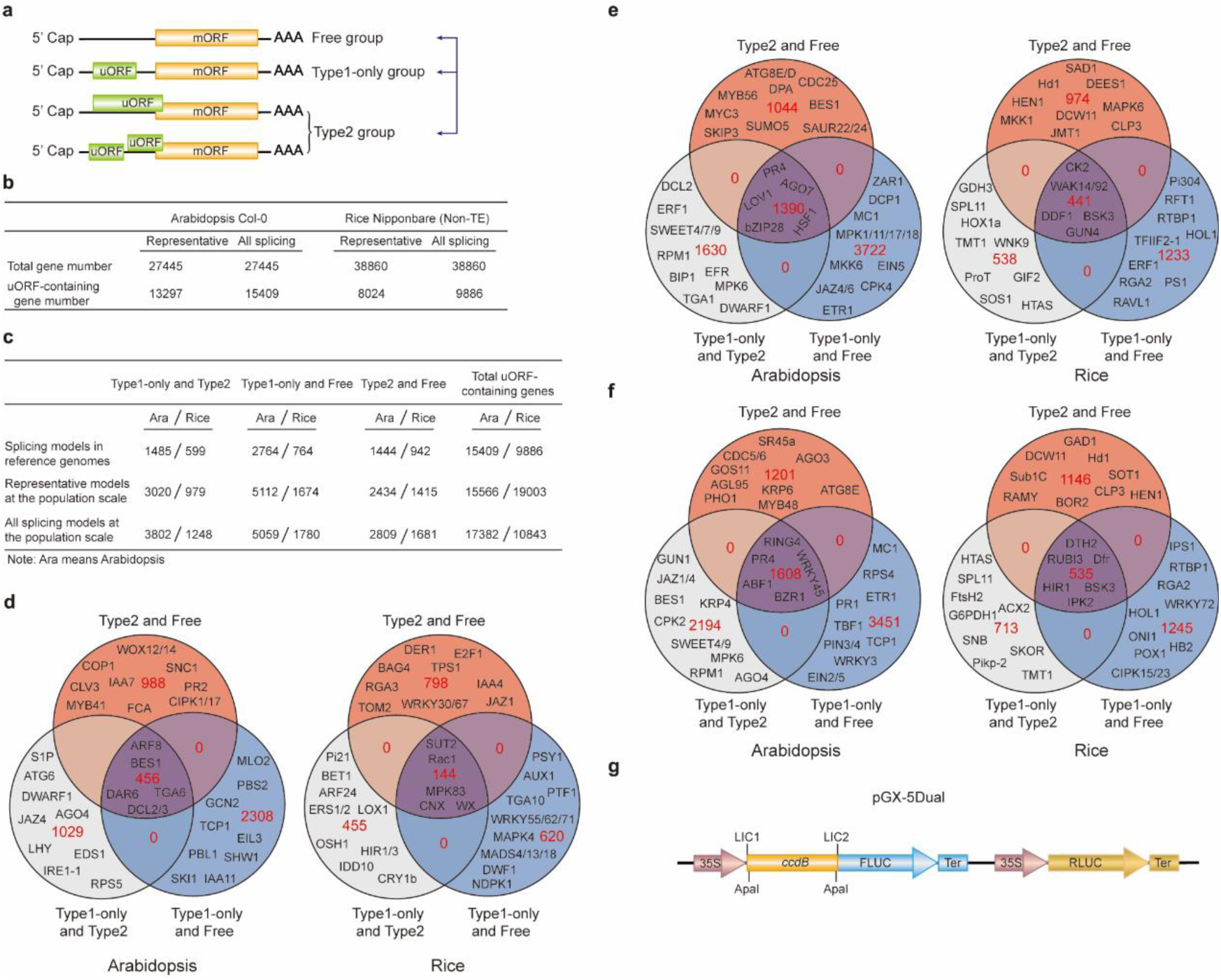
uORF variation. **a**, Schematic of altered uORF types analyzed in this study. Three Level1 uORF variants are focused by studying changes between Type1-only and Type2, between Type1-only and uORF-Free, and between Type2 and uORF-Free. Genes with both Type2 and Type1 on the same splicing model are grouped as Type2 in this analysis. **b**, Number of uORF-containing genes in Arabidopsis Col-0 and rice Nipponbare. The representative and all splicing gene models are calculated. Non-TE, none transposon element. **c**, Number of three Level1 uORF variants. Analysis is performed among different splicing models of reference genomes (Arabidopsis Col-0 and rice Nipponbare), and among accessions at the population scale using the representative and all splicing gene models. **d**, Venn diagram of three Level1 uORF variants within different splicing models of Arabidopsis Col-0 and rice Nipponbare. **e, f**, Venn diagram of three Level1 uORF variants of the representative gene models (e) and all splicing gene models at the population level (f). Example gene names are shown in different regions. **g**, Schematic of Dual-Luciferase vector pGX-5Dual. 5’ leader containing different uORF variants are cloned via LIC (ligation-independent cloning) method to control the translation of firefly luciferase (FLUC). ApaI is used to linearize the vector. Comparison of the ratio of FLUC activity and mRNA level to the internal control Renilla luciferase (RLUC) expressed from the same vector will indicate the effects of uORF variants on mRNA stability and translation efficiency.

In **uORF variation** menu, users are asked to input the gene model ID (e.g. AT4G36990) to search uORF variation among different splicing models in the reference genomes. Users can also input the splicing model ID (e.g. AT4G36990.1) and choose the accessions to return uORF variation in the selected population. The upper summary table lists all uORFs and the lower table returns uORFs listed according to accession names. The detailed information of uORF variants contains its frequency and the causal SNP or INDEL number at the popped out page by clicking ‘ALL’ in the summary table. Downloading at the top right corner allows the retrieval of all uORF information, including associated SNP or INDEL identifiers (VariationID). This information can also be accessed by clicking the individual accession name. SNP VariationID can be directly used to search their associated phenotypic variation using online tools such as https://aragwas.1001genomes.org for Arabidopsis (15), and http://snp-seek.irri.org/ and http://ricevarmap.ncpgr.cn/ for rice (23, 24, 28). These databases use different VariationID which can be converted using ID converter under the Tool menu.

The current GWAS databases use SNPs as genetic variants to perform association studies. Methods using INDELs are still under active development (33). On the other hand, many variables, such as population size, may cause GWAS false positive or biased associations (34). Considering uORF variants are responsible for 30%-80% protein abundance changes in human, we recommend performing a simple Dual-Luciferase assay to compare the translation efficiency of uORF variants by Agrobacterium-mediated transient expression in *Nicotiana benthamiana* (35, 36). We provide a Dual-Luciferase vector (pGX-5Dual) along with this database. Users can easily clone 5’ leaders containing different uORF alleles to the 5’ of firefly luciferase (FLUC) through ligation-independent cloning technology (37). Comparison of the ratio of FLUC activity and mRNA level to the internal control Renilla luciferase (RLUC) expressed from the same vector will indicate the effects of uORF variants on mRNA stability and translation efficiency. The difference will provide support for the association of uORF variation with its physiological roles deduced from its cognate mORF function (Figure 3g).

### CPuORF information

Analysis of evolutionary conservation suggests the existence of conserved peptide uORFs (CPuORFs) (38-44) and 97 non-redundant AUG-initiated CPuORFs in Arabidopsis have been confirmed here. It is suggested that most CPuORFs confer peptide-sequence dependent regulation in a cis manner, as metabolite receptors or sensors that function in the ribosome exit tunnel by stalling the ribosome and preventing reinitiation (4). There are also exceptional reports on uORFs functioning in trans-regulation (45, 46). In ‘uORF view’ menu, we collected all Arabidopsis genes containing CPuORF and also provided their rice homologues (Supplementary Table 5 in **Download** menu). This information will advance our understanding of uORF sequence-dependent regulation by facilitating the study of CPuORFs.

### Plant uORF reference curation

Eukaryotic uORF-related literatures to year 2013 have been curated in uORFdb through the Boolean search for key words in the NCBI PubMed database (47). To help the community get the latest view of the progress in plant uORF research, we manually curated all the relevant references on the basis of our knowledge and categorized the references into Case study, Mechanism study, Practical study, Genome-wide study, and Review. Users are invited to help us complete this section if missing or inappropriate references are found. Clicking the reference link will direct the users to the associated PubMed page or journal page.

### Customized uORF analysis

In the current database, we only processed uORFs with AUG as the initiation codon. With the rapid development of ribosome footprinting, mounting evidence suggests the usage of Non-AUG initiation codons in translational control (48). Those Non-AUG-initiated ORF-encoded peptides are also detectable by mass spectrometry (26). However, Non-AUG-initiated uORFs may function in an opposite way to AUG uORFs in translational control (48). This nuance remains poorly understood and systematic variation analysis of Non-AUG-initiated uORFs will be included when more information is available in plants. Therefore, users who are interested in Non-AUG studies or uORF studies in species other than Arabidopsis and rice are encouraged to use our searching tool under the menu of Tools to obtain the basic uORF information by inputting different uORF initiation codons and input cDNA and CDS sequences separately.

### Future Direction

uORFs are common in eukaryotes and information from more organisms will be useful additions to our database in the future. uORFs may encode functional peptides to act in either trans or cis manners, and this information will need to be evaluated by the combination of ribosome footprinting and mass-spectrometry data, which will be integrated as it becomes available. In addition, uORFs are RNA cis-elements which require trans-acting factors to regulate translation (36). Meanwhile, co-regulatory cis-elements, such as the R-motif identified in our previous study (36), may account for uORF regulation specificity and diversity. Information about regulatory trans-acting factors and co-acting cis-element variation will be integrated into the database progressively. Furthermore, a uORF calculator will be developed to predict the regulatory power of natural or synthetic uORFs for tailored protein expression after machine learning of large experimental data is achieved.

## Supplementary Information (in Download menu)

Supplementary Table 1. uORF information in Arabidopsis

Supplementary Table 2. uORF information in rice

Supplementary Table 3. uORF variation in Arabidopsis

Supplementary Table 4. uORF variation in rice

Supplementary Table 5. CPuORFs in Arabidopsis and rice

Supplementary Method: Ligation-independent cloning of Dual-LUC vector

## Acknowledgements

We thank Sophia Zebell and Paul J. Zwack at Duke University for comments.

